# Puberty timing and adiposity change across childhood and adolescence: disentangling cause and consequence

**DOI:** 10.1101/578005

**Authors:** Linda M O’Keeffe, Monika Frysz, Joshua A. Bell, Laura D. Howe, Abigail Fraser

**Author notes:** Denotes equal contribution (Laura D. Howe, Abigail Fraser). **Email addresses of authors:**. **Corresponding author:** Dr Linda M. O’Keeffe, MRC Integrative Epidemiology Unit at the University of Bristol, Oakfield House, Oakfield Grove, Bristol, UK, BS82BN.

## Abstract

**Objective:** To better understand if earlier puberty is more likely a result of adiposity gain in childhood than a cause of adiposity gain in adulthood.

**Design:** Prospective birth cohort study.

**Setting:** Population based study of children born in 1991/1992 in Bristol UK (Avon Longitudinal Study of Parents and Children (ALSPAC)).

**Participants:** 4,186 participants (2,176 female and 1,990 male) of predominantly White ethnicity with 18,232 repeated measures throughout follow-up.

**Exposures & outcomes:** Repeated measures of height from 5y to 20y to identify puberty timing (age at peak height velocity) and repeated measures of dual-energy X-ray absorptiometry-derived fat mass from age 9y to 18y, modelled separately in females and males using models based on chronological age and time before and after puberty onset.

**Results:** Mean age at peak height velocity was 11.7y (standard deviation (SD)=0.8y) for females and 13.6y (SD=0.9y) for males. In adjusted models of fat mass by chronological age, a one-year later age at peak height velocity was associated with 20.4% (95% Confidence Interval (CI): 18.5% to 22.3%) and 22.8% (95% (CI): 20.7% to 24.8%) lower fat mass in females and males respectively at 9y. These differences were smaller at age 18y: 7.8% (95% (CI):5.9% to 9.6%) and 11.9% (95% (CI): 9.1%, to 14.7%) lower fat mass in females and males respectively per year later age at peak height velocity. Trajectories of fat mass by time before and after puberty onset provided strong evidence for an association of pre-pubertal fat mass with puberty timing, and little evidence of an association of puberty timing with post-pubertal changes in fat mass in females. In males, findings were less clear before puberty though there was some evidence for an association of earlier puberty timing with great post-pubertal gain in fat mass.

**Conclusions:** Earlier puberty is more likely a result of adiposity gain in childhood than a cause of adiposity gain in adulthood in females. In males early to puberty, differences in fat mass after puberty are driven partially by tracking of adiposity from early childhood but also greater gains in post-pubertal adiposity. Reducing levels of childhood adiposity may help prevent both earlier puberty, later life adiposity and their associated adverse social, mental and physical health sequelae.

## Introduction

Age at puberty onset has decreased substantially among females since the mid-1900s ^1^ Secular trends in males are less well understood due to imprecise markers of pubertal age such as age at voice breaking compared with age at menarche among females ^1 2^. Earlier puberty directly results in younger fertility and thus carries important social implications ^3^, but it may also have adverse implications for health, with evidence of increased risk of adult obesity, type 2 diabetes, cardiovascular disease, and several cancers in both sexes ^4–7^.

Higher body mass index (BMI) before puberty onset is associated with earlier menarche in females, raising the possibility that much of the associations of puberty timing with health in later life reflects tracking of adiposity from childhood ^2 8–12^. In males, however, some studies have found that higher childhood BMI is associated with later puberty, ^10 13 14^, while others have found associations similar to those observed in females ^8 9 15–17^. In a recent systematic review and meta-analysis of 11 cohort studies, pre-pubertal obesity among females was associated with earlier menarche but there was insufficient and inconsistent evidence in males ^18^. A recent Mendelian randomisation (MR) analysis suggested that earlier age at menarche causes higher adult BMI but lacked data on pre-pubertal BMI for adjustment ^19^. In contrast, another recent MR which did have pre-pubertal BMI data suggested that associations of earlier puberty with higher adult BMI are largely confounded by childhood BMI ^20^. Thus, whether reducing childhood adiposity is important for the prevention of early puberty in both sexes and if prevention of early puberty also has additional benefits for the prevention of adiposity and its associated health outcomes in adulthood is unclear.

Most prospective studies to date have used self-reported measures of puberty timing such as age of voice breaking in males, have lacked data on pre and post-pubertal adiposity together and relied predominantly on indirect measures of adiposity such as BMI. In addition, disentangling direction of causality of puberty timing and adiposity may be difficult in available MR studies due to a shared genetic architecture between adiposity and puberty timing (age at menarche) ^7^. In this study, we aimed to better understand the association between puberty timing and pre-and post-pubertal adiposity change by examining an objective growth-based measure of pubertal onset (age at peak height velocity (aPHV)) in relation to change in directly measured dual-energy X-ray absorptiometry-derived (DXA-derived) fat mass across childhood and adolescence.

## Methods

### Study participants

Data were from the Avon Longitudinal Study of Parents and Children (ALSPAC), a prospective birth cohort study in southwest England ^21 22^. Pregnant women resident in one of the three Bristol-based health districts with an expected delivery date between April 1, 1991 and December 31, 1992 were invited to participate. The study is described elsewhere in detail ^21 22^. ALSPAC initially enrolled a cohort of 14,451 pregnancies, from which 13,867 live births occurred in 13,761 women. Follow-up has included parent- and child-completed questionnaires, clinic attendance, and links to routine data. Research clinics were held when the participants were approximately 7y, 9y, 10y, 11y, 13y, 15y, and 18y. Ethical approval for the study was obtained from the ALSPAC Ethics and Law Committee and the Local Research Ethics Committees. The study website contains details of all the data that is available through a fully searchable data dictionary http://www.bristol.ac.uk/alspac/researchers/our-data/^23^.

### Data

#### Assessment of puberty timing

Puberty is a period of intense hormonal activity and rapid growth, of which the most striking feature is the spurt in height ^24^. aPHV is a validated measure of pubertal timing ^24^ captured using Superimposition by Translation and Rotation (SITAR), a non-linear multilevel model with natural cubic splines which estimates the population average growth curve and departures from it as random effects ^25 26^. Using SITAR, PHV was identified in ALSPAC participants using numerical differentiation of the individually predicted growth curves, with aPHV being the age at which the maximum velocity is observed ^25–27^. Repeated height data included measurements from research clinics. Individuals with at least one measurement of height from 5y to <10y, 10y to < 15y and 15y to 20y are included here. Data was analysed for females and males separately. The model was fitted using the SITAR package in R version 3.4.1. Further details of height measures are included in eTable 1 and how aPHV was derived is described elsewhere ^27^ and in Supplementary Material.

#### Assessment of adiposity

Adiposity was assessed via total body fat mass (in kg, less head) as derived from whole body DXA scans performed five times at ages 9y, 11y, 13y, 15y, and 18y using a GE Lunar Prodigy (Madison, WI, USA) narrow fan beam densitometer.

#### Co-variates

We considered the following as potential confounders: birth weight, gestational age, maternal education, parity, maternal smoking during pregnancy, maternal age, maternal pre-pregnancy BMI, household social class, marital status, partner education and ever breastfeeding (all measured by mother- or mother’s partner-completed questionnaires; details in Supplementary Material).

### Statistical analysis

Multilevel models were used to examine change in fat mass during childhood and adolescence ^28 29^. Using terms such as polynomials and splines to account for non-linearity in the trajectory, such models can estimate mean trajectories of the outcome while accounting for the non-independence or clustering of repeated measurements within individuals, change in scale and variance of measures over time, and differences in the number and timing of measurements between individuals (using all available data from all eligible participants under a missing at-random assumption (MAR)) ^30 31^.

Participants who had a measure of aPHV, at least one measure of fat mass from 9y to 18y and complete data on all confounders were included in analyses, leading to a total sample of 4,176 (2,186 females and 1,990 males). Participants who reported being pregnant at the 18-year clinic were excluded from the multilevel models at that time point only (N=6). All models were adjusted for height using the time- and sex-varying power of height that best resulted in a height-invariant measure, described in detail elsewhere ^32^. Fat mass was log transformed prior to analysis due to its skewed distribution. All analyses were performed separately for females and males. aPHV was normally distributed in both sexes. Linearity of associations was examined by comparing the model fit of regressions of fat mass on aPHV, with aPHV treated as a continuous exposure and as a categorical exposure (thirds of aPHV). Model fit was formally tested using a likelihood ratio test. Prior to analysis, aPHV was centred on the sex-specific mean of aPHV for females and males. We performed unadjusted and confounder adjusted analyses for all models.

From all models, we back-transformed the difference in fat mass trajectories per year of aPHV and the average trajectory for the 10^th^, median and 90^th^ sex-specific percentile of aPHV. The back-transformed difference in fat mass per year of aPHV is a ratio of geometric means, expressed here as a percentage difference per year of aPHV. The average trajectories back-transformed from the log scale are presented in figures.

#### Models for fat mass trajectories

A common approach to modelling change over time using multilevel models involves examining change by chronological age ^33 34^. However, when change before or after a specified event is of interest (for example, onset of puberty or menopause), it is also possible to model change according to other time metrics such as time before and/or after the event. Thus, to gain a greater understanding of the association of aPHV with change in fat mass during childhood and adolescence, we modelled trajectories of fat mass in two ways: by chronological age, and separately by time before and after puberty.

#### Model 1: Chronological age-based models

Fat mass was previously modelled according to chronological age using linear spine multilevel models, with three periods of linear change (9-13y, 13-15y and 15-18y) ^32–35^. Thus, for this analysis, we examined whether this model was appropriate for modelling change over time within quartiles of pubertal age to ensure that model fit was good across the entire distribution of pubertal age. Subsequently, the association between aPHV and chronological age-based trajectories was then examined for females and males by including an interaction between centred sex-specific aPHV and the intercept (age 9y) and each spline period, providing an estimate of the difference in the average trajectory of fat mass from age 9y to 18y, per year later aPHV. Confounders were included as interactions with both the intercept and linear slopes.

#### Model 2: Pubertal age-based models

The purpose of the pubertal age-based model is to examine whether changes in fat mass before or after puberty onset differ by aPHV. In order to select an appropriate model, we examined observed data for fat mass in females and males by sex-specific quartiles of aPHV. Based on the observed data, a selection of suitable models was examined, each with different numbers of pre-and post-pubertal change periods. We compared observed and predicted values of fat mass for these models by sex-specific quartiles of pubertal age to examine model fit. In females, the final model selected for change in fat mass included two periods of change (pre-puberty and post-puberty). The final model for change in fat mass in males had three periods of change (from 9y to three years before puberty, from three years before puberty to puberty (i.e. aPHV) and from puberty to the end of follow-up at 18y). Differences in the rate of change in fat mass before and after puberty by aPHV were then modelled by including an interaction between centred sex-specific aPHV and the intercept (fat mass at puberty) and each linear spline period (one pre-and post-pubertal spline period for females and two pre-and one post-pubertal spline period for males). This model provided insight into whether different ages at PHV were accompanied by different rates of change in fat mass before and after puberty. Confounders were included as interactions with the intercept and linear slopes.

Details of model fit for both models are included in eTables 2 & 3.

#### Additional and sensitivity analyses

We examined the characteristics of mothers of participants included in our analysis compared with mothers of participants excluded from our analysis due to missing exposure, outcome or confounder data, using maternal socio-demographic characteristics measured at or close to birth to better understand generalisability and the potential for selection bias. We regressed observed fat mass at 9y (first available measure) and 18y (last occasion of measurement) on aPHV in females and males and compared results to those obtained from the multilevel models at these ages. We performed sensitivity analyses restricting the sample to participants with at least one fat mass measure before and one after aPHV, to examine whether results from the main analysis were driven by participants with only a single pre- or post-puberty fat mass measure. In addition, we performed a sensitivity analysis among females examining whether the association of self-reported age at menarche with fat mass during childhood and adolescence was similar to findings for the association of aPHV and fat mass.

## Results

The characteristics of participants included in analyses, by sex, are shown in Table 1. Mean aPHV was 11.7 (standard deviation (SD) = 0.8) for females (N=2,176) and 13.6 (SD=0.9) for males (N=1,990). Findings from linearity tests of aPHV and fat mass at each age found little evidence of departure from linearity, allowing aPHV to be examined as a continuous exposure (see eTable 4). Mothers of participants included in the analysis were more likely to be married, have higher household social class, higher education, higher partner education, lower prevalence of smoking during pregnancy, lower parity and higher maternal age compared with participants excluded due to missing exposure, outcome or confounder data (eTable 5). Participants included in analyses were more likely to be female, have higher gestational age at birth and higher birth weight compared with those excluded. However, maternal BMI and aPHV were similar between included and excluded participants (eTable 5).

**Table 1.** Characteristics of ALSPAC participants included in the analysis, by sex

|  | Females<br>N=2,176 | Males<br>N=1,990 | P value for comparison<sup>a</sup> |
| --- | --- | --- | --- |
|  | n (%) | n (%) |  |
| **Maternal marital status** |  |  |  |
| Never married | 269(12.3) | 196(9.8) | <0.0001 |
| Widowed | 1(0.0) | 3(0.2) |  |
| Divorced | 65(3.0) | 70(3.5) |  |
| Separated | 20(0.9) | 15(0.8) |  |
| 1<sup>st</sup> Marriage | 1701(77.8) | 1564(78.6) |  |
| Marriage 2 or 3 | 130(5.9) | 142(7.1) |  |
| **Household social class<sup>b</sup>** |  |  |  |
| Professional | 377(17.2) | 390(19.6) | <0.0001 |
| Managerial & Technical | 1009(46.2) | 947(47.6) |  |
| Non-Manual | 526(24.1) | 449(22.6) |  |
| Manual | 198(9.1) | 142(7.1) |  |
| Part Skilled & Unskilled | 76(3.5) | 62(3.1) |  |
| **Maternal education** |  |  |  |
| Less than O level | 360(16.5) | 311(15.6) | <0.05 |
| O level | 781(35.7) | 694(34.9) |  |
| A level | 621(28.4) | 602(30.3) |  |
| Degree or above | 424(19.4) | 383(19.2) |  |
| **Mother's Partner's highest educational qualification** |  |  |  |
| Less than O level | 553(25.3) | 426(21.4) | <0.0001 |
| O level | 471(21.5) | 444(22.3) |  |
| A level | 640(29.3) | 583(29.3) |  |
| Degree or Above | 522(23.9) | 537(27.0) |  |
| **Maternal smoking during pregnancy** |  |  |  |
| No | 1874(85.7) | 1707(85.8) | 0.79 |
| Yes | 312(14.3) | 283(14.2) |  |
| **Parity** |  |  |  |
| 0 | 1068(48.9) | 981(49.3) | <0.0001 |
| 1 | 796(36.4) | 684(34.4) |  |
| 2 | 322(14.7) | 325(16.3) |  |
| **Breastfeeding** |  |  |  |
| Exclusive | 841(38.5) | 697(35.0) | <0.01 |
| Non-exclusive | 1006(46.0) | 1034(52.0) |  |
| Never | 339(15.5) | 259(13.0) |  |
|  | **Mean (SD)** | **Mean (SD)** |  |
| **Gestational age (weeks)** | 39.6(1.6) | 39.4 (1.9) | <0.001 |
| **Birth weight (g)** | 3393.0(483.6) | 3481.4 (574.5) | <0.0001 |
| **Maternal pre-pregnancy BMI (kg/m<sup>2</sup>)** | 22.8(3.6) | 22.9(3.7) | 0.1 |
| **Maternal age at delivery (years)** | 29.3(4.3) | 29.7(4.4) | <0.001 |

<sup>a</sup> p value is for the difference in proportions for categorical variables from Chi squared test.
<sup>b</sup> Household social class was measured as the highest of the mother's or her partner's occupational social class using data on job title and details of occupation collected about the mother and her partner from the mother's questionnaire at 32 weeks gestation. Social class was derived using the standard occupational classification
(SOC) codes developed by the United Kingdom Office of Population Census and Surveys and classified as I professional, II managerial and technical, III NM non-manual, III M manual, and IV&V part skilled occupations and unskilled occupations. .

### Puberty timing and adiposity change from models by chronological age

A one-year later aPHV in females was associated with a lower fat mass at 9y (Table 2) and faster gain in fat mass from 9y to 18y. By 18y, the mean difference per year later aPHV persisted but was smaller. In males, associations were comparable; a one-year later aPHV was associated with lower fat mass at 9y which reduced to a smaller difference at age 18 (Table 2). Mean adjusted trajectories of fat mass from 9y to 18y for the 10^th^ (age 11 in females and age 13 males), 50^th (^age 12 in females and age 14 in males) and 90^th^ (age 13 in females and age 15 in males) sex-specific percentiles of aPHV by chronological age and are presented in Figure 1.

**Table 2.**
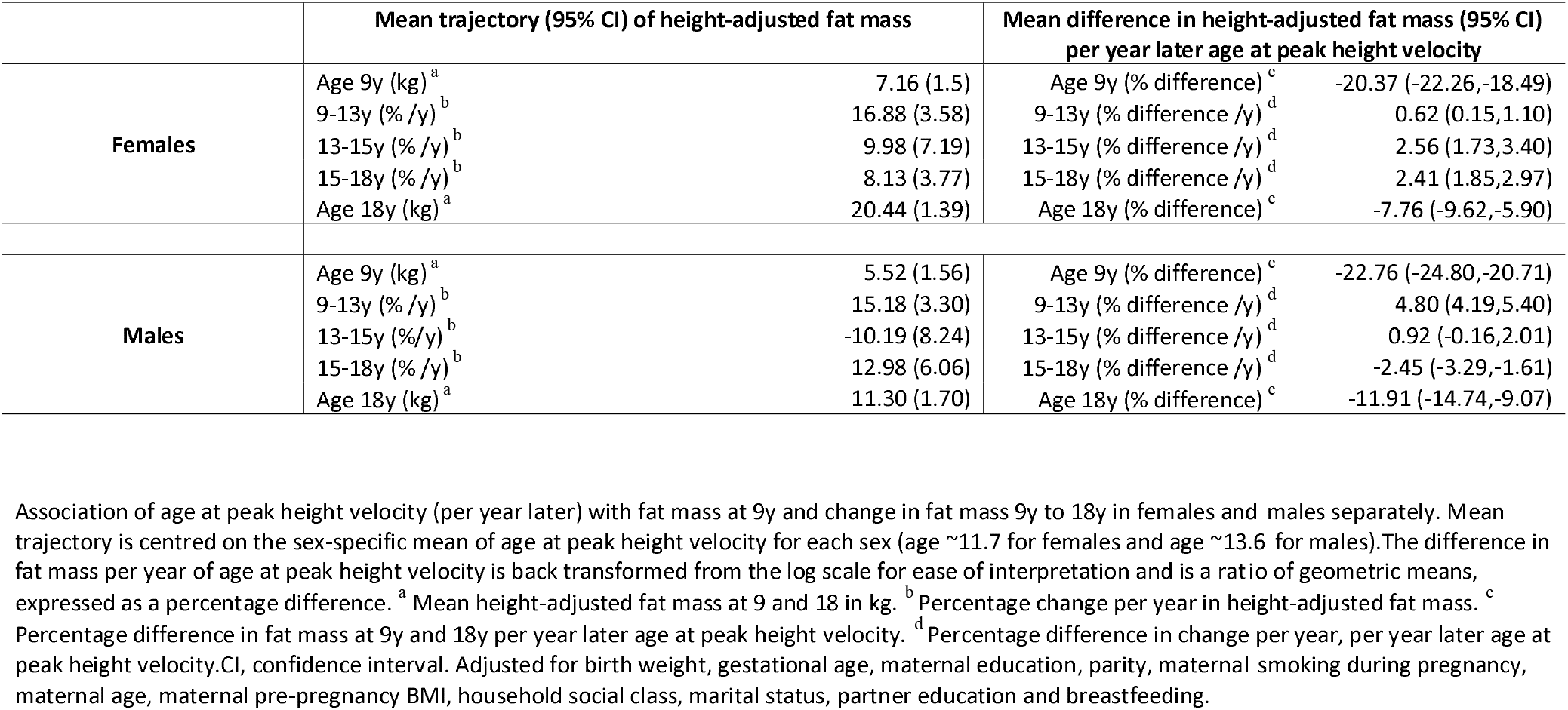
Adjusted mean trajectory and mean difference in trajectory of height-adjusted fat mass per year later age at peak heigh t velocity, from chronological age multilevel models

**Table 3.** Adjusted mean trajectory and mean difference in trajectory of height-adjusted fat mass per year later age at peak heigh t velocity, from pubertal age multilevel models

|  | Mean trajectory (SD) of height-adjusted fat mass |  | Mean difference in height-adjusted fat mass (95% CI)<br>per year later age at peak height velocity |  |
| --- | --- | --- | --- | --- |
| **Females** | Before puberty (% /y) <sup>a</sup> | 17.91 (2.35) | Before puberty (% difference /y) <sup>c</sup> | -1.38 (-2.15,-0.62) |
|  | Fat mass at puberty (kg) <sup>b</sup> | 11.25 (1.46) | Fat mass at puberty (% difference) <sup>d</sup> | -8.85 (-10.82,-6.88) |
|  | After puberty (%/y) <sup>a</sup> | 10.41 (3.57) | After puberty (% difference /y) <sup>c</sup> | 1.43 (1.10,1.76) |
| **Males** | Up-to 3 years before puberty (% /y) <sup>a</sup> | 22.50 (2.99) | Up-to 3 years before puberty (% difference/y) <sup>c</sup> | -5.87 (-7.22,-4.53) |
|  | From 3 years before to puberty (% /y) <sup>a</sup> | -0.12 (5.30) | From 3 years before to puberty (% difference/y) <sup>c</sup> | 3.47 (2.58,4.37) |
|  | Fat mass at puberty (kg) <sup>b</sup> | 8.05 (1.70) | Fat mass at puberty (% difference) <sup>d</sup> | -2.31 (-5.16,0.54) |
|  | After puberty (% /y ) <sup>a</sup> | 4.65 (4.00) | After puberty (% difference /y ) <sup>c</sup> | -0.79 (-1.41,-0.18) |

Association of age at peak height velocity (per year later) with change in fat mass before puberty, fat mass at puberty and change in fat mass after puberty in females and males separately. Mean trajectory is centred on the sex-specific mean of age at peak height velocity for each sex (age ~11.7 for females and age ~13.6 for males). The difference in fat mass per year of age at peak height velocity is back transformed from the log scale for ease of interpretation and is a ratio of geometric means, expressed as a percentage difference. <sup>a</sup> Mean height-adjusted fat mass at 9 and 18 in kg. <sup>b</sup> Percentage change per year in height-adjusted fat mass. <sup>c</sup> Percentage difference in fat mass at 9y and 18y per year later age at peak height velocity. <sup>d</sup> Percentage difference in change per year, per year later age at peak height velocity. CI, confidence interval. Models are adjusted for birth weight, gestational age, maternal education, parity, maternal smoking during pregnancy, maternal age, maternal pre-pregnancy BMI, household social class, marital status, partner education and breastfeeding.

**Figure 1.**
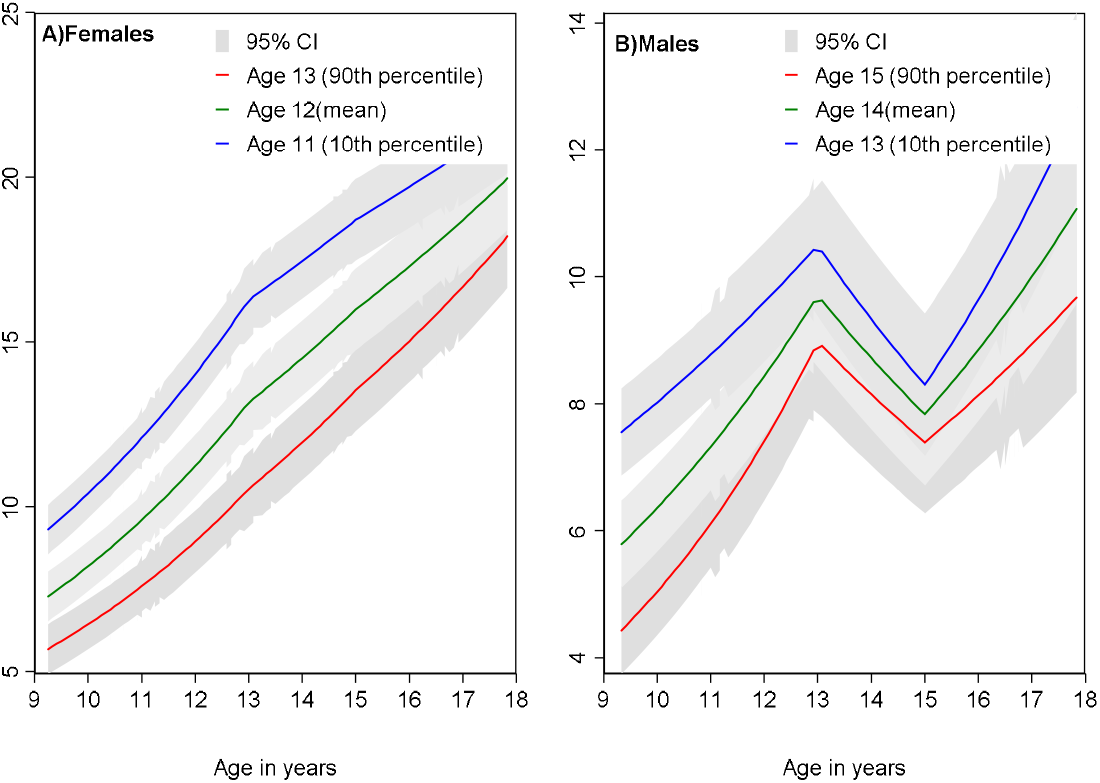
Mean adjusted trajectories of fat mass in females and males from 9y to 18y for the 10^th^, median and 90^th^ sex-specific percentiles of age at peak height velocity from multilevel models based on chronological age. Ages presented are rounded for ease of interpretation. Exact ages are 12.9y, 11.7y and 10.7y for females and 14.7y, 13.6y and 12.5y for males. Age at peak height velocity is normally distributed and median is equal to mean. Models are adjusted for birth weight, gestational age, mat ernal education, parity, maternal smoking during pregnancy, maternal age, maternal pre-pregnancy BMI, household social class, marital status, partner ed ucation and breastfeeding.

### Puberty timing and adiposity change from models by pubertal age

Among females, a one-year later aPHV was associated with a slower gain in fat mass before puberty, a lower fat mass at puberty and faster gain in fat mass after puberty. In males, up-to three years before puberty later aPHV was associated with slower gains in fat mass, whereas from three years before puberty to pubertal onset, later aPHV was associated with faster gain in fat mass. At puberty onset, males later to puberty had lower fat mass, albeit with confidence intervals spanning the null value. After puberty, later aPHV was associated with slower gain in fat mass. Mean adjusted trajectories of fat mass from 9y to 18y for the 10^th^ (age 11 in females and 13 males), 50^th^ (age 12 in females and 14 in males) and 90^th^ (age 13 in females and 15 in males) sex-specific percentiles of aPHV by pubertal age are presented in Figure 2.

**Figure 2.**
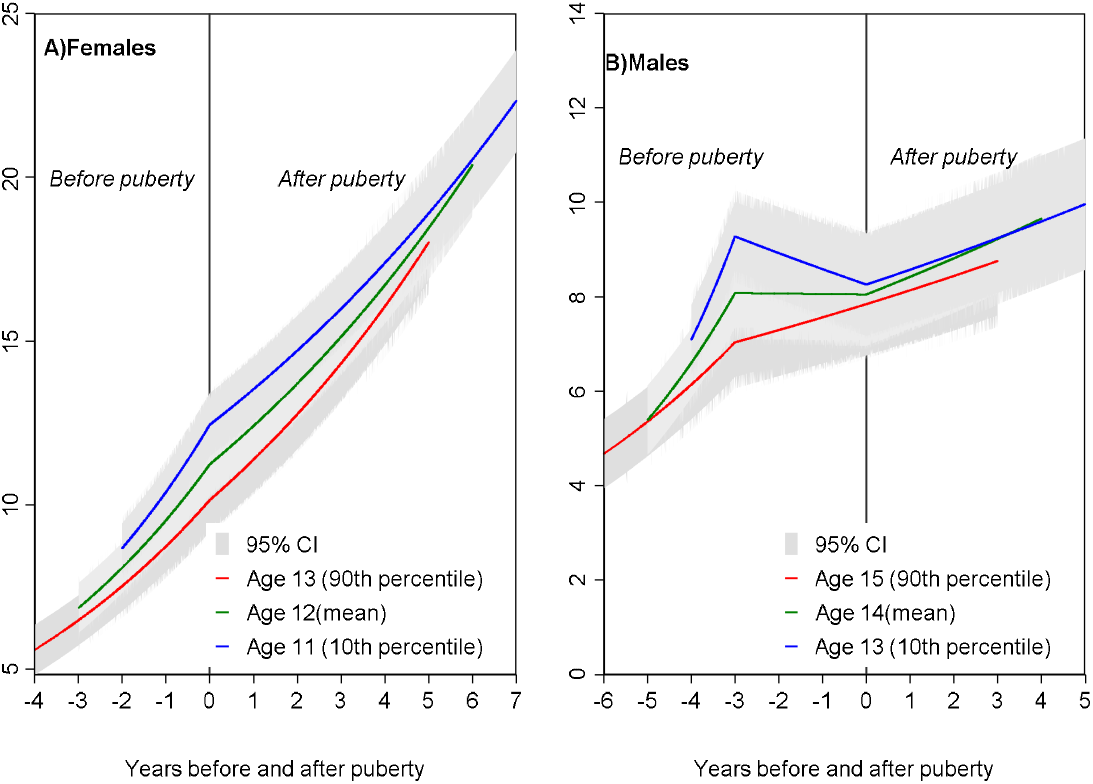
Mean adjusted trajectories of fat mass in females and males from 9y to 18y for the 10^th^, median and 90^th^ sex-specific percentiles of age at peak height velocity from multilevel models based on pubertal age. Ages presented are rounded for ease of interpretation. Exact ages are 12.9y, 11.7y and 10.7y for females and 14.7y, 13.6y and 12.5y for males. Age at peak height velocity is normally distributed and median is equal to mean. Models are adjusted for birth weight, gestational age, mat ernal education, parity, maternal smoking during pregnancy, maternal age, maternal pre-pregnancy BMI, household social class, marital status, partner ed ucation and breastfeeding.

Unadjusted and confounder-adjusted results were similar for each analysis (eTable 6 & 7).

### Sensitivity analyses

Estimates of sex-specific associations of aPHV with observed fat mass data at age 9y and 18y were similar to those obtained from both types of multilevel models (by chronological age and time before and after puberty onset) (eTable 8). Results were not appreciably different when analyses were restricted to participants with at least one measure before and one measure after aPHV (eFigure 2 & 3). Results for females were also similar when analysed using self-reported age at menarche rather than aPHV (eFigure 4).

## Discussion

This study aimed to better understand the nature of puberty timing and adiposity change by examining associations of an objective height-based measure of puberty timing (aPHV) with change in DXA-measured total body fat mass repeatedly measured throughout childhood and adolescence. In females, our findings suggest that earlier puberty timing is more likely to be the result of adiposity gain in childhood than a cause of adiposity gain in adulthood. In males, findings suggest that childhood adiposity may also contribute to early puberty timing and that differences in fat mass after puberty are driven partially by tracking of adiposity from early childhood but also greater gains in post-pubertal adiposity in males early to puberty. Altogether the findings suggest that reducing levels of childhood adiposity may help to prevent earlier puberty, adult adiposity and its adverse health and social outcomes.

### Comparison with other studies

Our findings in females are consistent with several previous studies showing inverse associations of pre-pubertal BMI and puberty timing ^2 8–10^. For example, in a Swedish study (N=3,650) which used growth data from age 7y to 18y years to estimate aPHV, faster rates of gain in BMI from 2y to 8y were associated with earlier puberty in females ^15^. In a Danish study (N=156, 835), BMI at 7y was associated with earlier puberty in females, based on onset of growth spurt and aPHV ^16^. These findings are also comparable to a report from the Cardiovascular Risk in Young Finns study (N=794) which concluded that greater childhood BMI contributed to earlier age at menarche and because of tracking, to greater adult BMI ^12^. Our results are also similar to findings from a recent MR in our cohort which showed that associations of age at menarche with adulthood BMI result from tracking of childhood BMI ^20^. Our findings showed greater gains in fat mass after puberty among females later to puberty which to our knowledge has not been demonstrated previously due to a lack of studies with repeated measures of pre-and post-pubertal fat mass. Though it is difficult to understand the precise reason for this, one possibility includes catch up in fat mass accrual after puberty among females later to puberty who have had slower rates of gain in fat mass prior to puberty.

Our results in males are consistent with most ^8 9 15–17^ but not all ^10 13^ previous studies that found that greater adiposity in early childhood is associated with early puberty. The Christ’s Hospital Cohort (N=1,520) showed that males with higher childhood BMI before puberty had earlier aPHV ^17^ while in the aforementioned Swedish study, change in BMI from 2y to 8y was also associated with earlier puberty ^15^. Similarly, a Swedish study of 99 monozygotic and 76 dizygotic twins found that early childhood BMI was associated with earlier puberty in males. ^9^ One known exception to this is a US study of 401 males showing that faster gains in BMI from two to 11.5y were associated with later puberty onset in males, based on Tanner staging as assessed by paediatric endocrinologists ^13^. Our findings of greater gains in fat mass up to three years before puberty followed by slower gains in fat mass in the period directly before puberty among males early to puberty build on and partially consolidate these inconsistent findings to date in males. Increasing body fat is thought to play a critical role in switching on adrenal androgen secretion leading to the initiation of puberty; this may explain steeper rises in fat mass in males earlier to puberty up to three years before puberty ^2^. Once the underlying process of puberty is initiated in males, fat mass decreases and this decrease is steeper in males with a younger age at puberty. These decreases in fat mass in males prior to puberty may be linked to rising testosterone levels in males ^2^. aPHV has also been shown to be a marker of more advanced puberty stages in males than in females (Tanner genetalia stages 4 and 5 in males compared with Tanner breast stages 2 and 3 in females)^36^. These differences may contribute to the different associations between fat mass change and a PHV observed for males and females as well as to the contrasting associations of aPHV with fat mass change up to 3 years before puberty and then between 3 years and puberty and aPHV.

### Strengths and limitations

The main strengths of our study include the use of an objective measure of puberty timing (aPHV) based on prospective, repeated measures of height from age 5y to 20y which is a more accurate marker than age of voice breaking or Tanner staging, measures that have been frequently used in previous studies. We also used repeated measures of adiposity from before to after puberty onset which were directly measured using DXA scans and have not been available in previous studies. Limitations include the lack of measures of fat mass before 9y and the availability of few measures around puberty, which limit our ability to detect subtle and/or acute changes in fat mass around puberty. Missing data due to loss to follow-up was associated with greater social disadvantage. However, we aimed to minimise potential bias by including all participants with at least one measure of fat mass from age 9y to 18y.

## Conclusion

Earlier puberty timing is likely to be more of a result of adiposity gain in childhood than a cause of adiposity gain in adulthood in females. In males, differences in fat mass after puberty are driven partially by tracking of adiposity from early childhood but also by greater gains in post-pubertal adiposity in males earlier to puberty compared with males later to puberty. Reducing levels of childhood adiposity may help prevent earlier puberty, adult adiposity and their adverse health and social outcomes.

## Contributor and guarantor statement

LMOK had the idea for the study, performed all analyses and wrote the manuscript up for publication. MF derived age at peak height velocity. MF, JB, AF and LDH interpreted the results and provided critical revisions to the manuscript. LMOK will act as a guarantor for the work.

## Competing interest’s declaration

None of the authors have any conflicts of interest to declare.

## Public and patient involvement

Public and patient involvement was not part of this research.

## Sources of funding and role of funding source

The UK Medical Research Council and Wellcome (Grant ref: 102215/2/13/2) and the University of Bristol provide core support for ALSPAC. This publication is the work of the authors and LMOK will serve as guarantor for the contents of this paper. LMOK is supported by a UK Medical Research Council Population Health Scientist fellowship (MR/M014509/1). LDH and AF are supported by Career Development Awards from the UK Medical Research Council (grants MR/M020894/1 and MR/M009351/1, respectively). All authors work in a unit that receives funds from the UK Medical Research Council (grant MC_UU_00011/3, MC_UU_00011/6). These funding sources had no role in the design and conduct of this study.

## Data sharing

Data are available upon submission and approval of a research proposal to the ALSPAC Executive. Further information can be found at http://www.bristol.ac.uk/alspac/researchers/access/

## Acknowledgements

We are extremely grateful to all the families who took part in this study, the midwives for their help in recruiting them, and the whole ALSPAC team, which includes interviewers, computer and laboratory technicians, clerical workers, research scientists, volunteers, managers, receptionists and nurses.

## References

1. Euling SY, Herman-Giddens ME, Lee PA, et al. Examination of US puberty-timing data from 1940 to 1994 for secular trends: panel findings. Pediatrics 2008;121(Supplement 3):S172–S91.

2. Kaplowitz PB. Link between body fat and the timing of puberty. Pediatrics 2008;121(Supplement 3):S208–S17.

3. Golub MS, Collman GW, Foster PM, et al. Public health implications of altered puberty timing. Pediatrics International 2008;121(Supplement 3):S218–S30.

4. Prentice P, Viner RM. Pubertal timing and adult obesity and cardiometabolic risk in women and men: a systematic review and meta-analysis. International Journal of Obesity 2013;37(8):1036.

5. Canoy D, Beral V, Balkwill A, et al. Age at menarche and risks of coronary heart and other vascular diseases in a large UK cohort. Circulation 2014:CIRCULATIONAHA. 114.010070.

6. Charalampopoulos D, McLoughlin A, Elks CE, et al. Age at menarche and risks of all-cause and cardiovascular death: a systematic review and meta-analysis. American journal of epidemiology 2014;180(1):29–40.

7. Day FR, Elks CE, Murray A, et al. Puberty timing associated with diabetes, cardiovascular disease and also diverse health outcomes in men and women: the UK Biobank study. Scientific Reports 2015;5:11208.

8. Buyken AE, Karaolis-Danckert N, Remer T. Association of prepubertal body composition in healthy girls and boys with the timing of early and late pubertal markers. The American journal of clinical nutrition 2008;89(1):221–30.

9. Silventoinen K, Haukka J, Dunkel L, et al. Genetics of pubertal timing and its associations with relative weight in childhood and adult height: the Swedish Young Male Twins Study. Pediatrics 2008;121(4): e885–e91.

10. Wang Y. Is obesity associated with early sexual maturation? A comparison of the association in American boys versus girls. Pediatrics 2002;110(5):903–10.

11. Lee JM, Appugliese D, Kaciroti N, et al. Weight status in young girls and the onset of puberty. Pediatrics 2007;119(3):e624–e30.

12. Kivimèki M, Lawlor DA, Smith GD, et al. Association of age at menarche with cardiovascular risk factors, vascular structure, and function in adulthood: the Cardiovascular Risk in Young Finns study. The American journal of clinical nutrition 2008;87(6):1876–82.

13. Lee JM, Kaciroti N, Appugliese D, et al. Body mass index and timing of pubertal initiation in boys. Archives of pediatrics & adolescent medicine 2010;164(2):139–44.

14. Lee JM, Wasserman R, Kaciroti N, et al. Timing of puberty in overweight versus obese boys. Pediatrics 2016:peds. 2015–0164.

15. He Q, Karlberg J. BMI in childhood and its association with height gain, timing of puberty, and final height. Pediatric Research 2001;49(2):244.

16. Aksglaede L, Juul A, Olsen LW, et al. Age at puberty and the emerging obesity epidemic. PloS one 2009;4(12):e8450.

17. Sandhu J, Ben-Shlomo Y, Cole TJ, et al. The impact of childhood body mass index on timing of puberty, adult stature and obesity: a follow-up study based on adolescent anthropometry recorded at Christ’s Hospital (1936-1964). International Journal of Obesity 2006;30(1):14.

18. Li W, Liu Q, Deng X, et al. Association between Obesity and Puberty Timing: A Systematic Review and Meta-Analysis. International Journal of Environmental Research and Public Health 2017;14(10):1266.

19. Gill D, Brewer CF, Fabiola Del Greco M, et al. Age at menarche and adult body mass index: a Mendelian randomization study. International Journal of Obesity 2018:1.

20. Bell JA, Carslake D, Wade KH, et al. Influence of puberty timing on adiposity and cardiometabolic traits: A Mendelian randomisation study. PLoS medicine 2018;15(8):e1002641.

21. Boyd A, Golding J, Macleod J, et al. Cohort Profile: The ‘Children of the 90s’—the index offspring of the Avon Longitudinal Study of Parents and Children. International journal of epidemiology 2013;42(1):111–27.

22. Fraser A, Macdonald-Wallis C, Tilling K, et al. Cohort profile: the Avon Longitudinal Study of Parents and Children: ALSPAC mothers cohort. International journal of epidemiology 2013;42(1):97–110.

23. University of Bristol. Avon Longitudinal Study of Parents and Children 2017 [Available from: http://www.bristol.ac.uk/alspac/researchers/access/.

24. Cole T, Pan H, Butler G. A mixed effects model to estimate timing and intensity of pubertal growth from height and secondary sexual characteristics. Annals of Human Biology 2014;41(1):76–83.

25. Cole TJ, Donaldson MD, Ben-Shlomo Y. SITAR--a useful instrument for growth curve analysis. International journal of epidemiology 2010;39(6):1558–66. doi: 10.1093/ije/dyq115 [published Online First: 2010/07/22]

26. Simpkin AJ, Sayers A, Gilthorpe MS, et al. Modelling height in adolescence: a comparison of methods for estimating the age at peak height velocity. Annals of human biology 2017;44(8):715–22.

27. Frysz M, Howe LD, Tobias JH, et al. Using SITAR (SuperImposition by Translation and Rotation) to estimate age at peak height velocity in Avon Longitudinal Study of Parents and Children. Wellcome Open Research 2018;3

28. Goldstein H. Multilevel statistical models; 2nd edition ed. London:: Edward Arnold 1995.

29. Laird NM, Ware JH. Random-effects models for longitudinal data. Biometrics 1982:963–74.

30. Howe LD, Tilling K, Matijasevich A, et al. Linear spline multilevel models for summarising childhood growth trajectories: A guide to their application using examples from five birth cohorts. Statistical Methods in Medical Research 2013:0962280213503925.

31. Tilling K, Macdonald-Wallis C, Lawlor DA, et al. Modelling childhood growth using fractional polynomials and linear splines. Annals of Nutrition and Metabolism 2014;65(2-3):129–38.

32. O’Keeffe L, Simpkin A, Tilling K, et al. Data on trajectories of measures of cardiovascular health in the Avon Longitudinal Study of Parents and Children (ALSPAC). Data in Brief 2018 33.

33. O’Keeffe L, Simpkin A, Tilling K, et al. Sex-specific trajectories of cardiometabolic risk factors during childhood and adolescence: a prospective cohort study Atherosclerosis. Atherosclerosis 2018;278:190–96.

34. O’Keeffe LM, Howe LD, Fraser A, et al. Associations of Y chromosomal haplogroups with cardiometabolic risk factors and subclinical vascular measures in males during childhood and adolescence. Atherosclerosis 2018;274:94–103.

35. Howe LD, Parmar PG, Paternoster L, et al. Genetic Influences on Trajectories of Systolic Blood Pressure Across Childhood and Adolescence. Circulation: Cardiovascular Genetics 2013;6(6):608–14.

36. Granados A, Gebremariam A, Lee JM. Relationship between timing of peak height velocity and pubertal staging in boys and girls. Journal of Clinical Research in Pediatric Endocrinology 2015;7(3):235.

