## Supplemental Material for "Puberty timing and adiposity change across childhood and adolescence: disentangling cause and consequence"

**Appendix 1: Details of deriving age at peak height velocity**

*Height measures included in the analysis*

Height data from questionnaires and health records were excluded. Only data measured at clinic assessments carried out after age 5y were included in the analysis. A Child in Focus (CIF) clinic measured height using a Leicester Height Measure on a 10% sub-sample of participants measured at age five. From 7y onwards, standing height was measured to the last complete mm using the Harpenden Stadiometer at clinics carried out at ages 7y, 9y, 10y, 11y, 13y, 14y, 15y and 18y. Data were further restricted to only include individuals with at least one measurement of height from 5y to <10y, 10y to < 15y and 15y to 20y. The final dataset for analysis included 20,849 height measurements for 2,688 boys and 24,216 measurements for 3,019 girls. The number of height measures for females and males is shown in Table 1. Further details of how age at peak height velocity (aPHV) was derived are described elsewhere ^1^.

**Table 1 Details of height measures available for deriving aPHV**

| **Males** |  |  | **Females** |  |  |
| --- | --- | --- | --- | --- | --- |
| **No. of height measurements per individual** | **Number of individuals** | **%** | **No. of height measurements per individual** | **Number of individuals** | **%** |
| 1 | 8 | 0.3 | N/A |  |  |
| 2 | 24 | 0.89 | 2 | 18 | 0.6 |
| 3 | 26 | 0.97 | 3 | 31 | 1.03 |
| 4 | 70 | 2.6 | 4 | 60 | 1.99 |
| 5 | 120 | 4.5 | 5 | 100 | 3.31 |
| 6 | 219 | 8.2 | 6 | 192 | 6.36 |
| 7 | 432 | 16.1 | 7 | 355 | 11.8 |
| 8 | 638 | 23.7 | 8 | 826 | 27.4 |
| 9 | 1117 | 41.6 | 9 | 1268 | 42 |
| 10 | 34 | 1.26 | 10 | 169 | 5.6 |
| Total | 2688 | 100% | Total | 3019 | 100% |

**Analysis deriving aPHV**

Available height measures were analysed for females and males separately using Superimposition by Translation and Rotation (SITAR) growth curve analysis with five degrees of freedom ^2^. This method is a validated method of deriving aPHV and is described elsewhere in detail ^2 3^. aPHV was defined as the age when the first derivative of the mean curve, plotted as height versus age, was maximal. After fitting the initial model, the data were checked and points with velocity exceeding four standard deviations (SDs) and standardized residuals exceeding three in absolute value were removed. The model explained 98.5% of variance in males and 98.7% in females. Mean growth curve and velocity plots for females and males are shown in Figure 1.

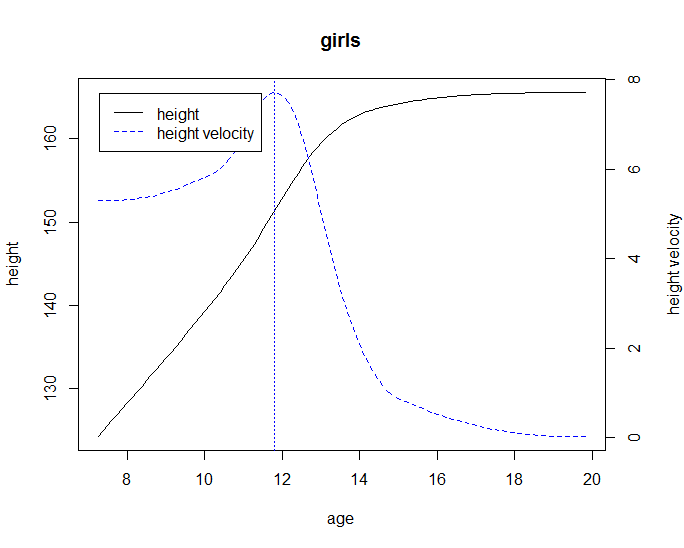

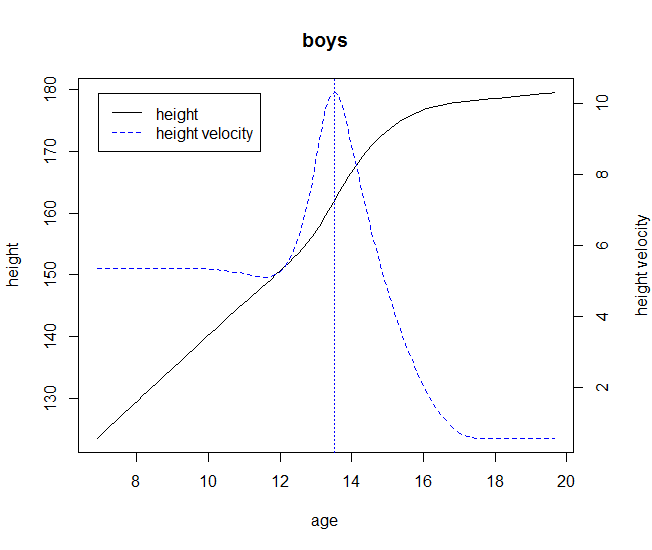

**Figure 1 Mean growth curve (black line) and velocity (blue dashed-line) plots estimated by SITAR for females and males.**

Vertical dotted line represents age at peak height velocity.

**Appendix 2 Details of model selection**

Fat mass was measured on five occasions between 9y and 18y. Values of fat mass four SDs greater than or less than the mean were excluded from the analysis. Fat mass was log transformed due to skewness of the data. We included all participants with at least one measure of fat mass in each multilevel model, under a missing at random (MAR), to minimise selection bias. The observations of participants who reported being pregnant at the 18-year clinic were excluded from the multilevel models at that time point only. Trajectories were modelled separately for females and males to allow each sex to have different variance-covariance matrices. Models were adjusted for a time-and sex-varying height co-variate which was included as a fixed effect, as described elsewhere in detail.

We modelled sex-specific change over time in fat mass according to chronological age and pubertal age to better understand the association of age at peak height velocity with change in fat mass during childhood and adolescence. In both models, linear splines were used to examine change in fat mass ^4^.

**Models based on chronological age**

A chronological age model was developed previously and is described elsewhere in detail ^4 5^ . Thus, for this analysis, we re-examined the fit of this model according to sex-specific quartiles of pubertal age to ensure that model fit was similar across each of the quartiles of pubertal age and that modifications to this model (including different periods of change or fewer or greater spline periods were not required) for different pubertal age groups. We found that this model had good model fit across sex-specific quartiles of pubertal age. In brief, age in years was centred at 9y. Knots were placed at 13y and 15y resulting in three periods of change; from 9-13y, 13-15y, 15-18y. As each sex was modelled separately, the models for males and females took the form of: log fat mass_ij_ = β_0_ + u_0j_ + (β_1_+ u_1j_ )s_ij1_ + (β_2_+ u_2j_ )s_ij2_ + (β_3_ + u_3j )_s_ij3_ + β_4_ (age&sex adjusted height covariate)_ij_ + e_ij_ where for person j at measurement occasion i; β_0_ represents the fixed effect coefficient for the average intercept, β_1_ to β_3_ represent fixed effect coefficients for the average linear slopes of each linear spline, β_4_ represents the fixed effect coefficient for the average difference in measurements between individuals of different heights, u_0j_ to u_3j_ indicate person-specific random effects for the intercept and slopes respectively, and e_ij_ represents the occasion-specific residuals or measurement error which was allowed to vary with age.

**Models based on pubertal age**

Models examining change in fat mass according to pubertal age were modelled de novo for this paper. We examined observed data at each age by sex to examine whether the shape of change over time was similar or different between quartiles of pubertal age. We found that the shape of change over time was similar across quartiles of pubertal age in each sex, but that change over time differed for females and males. Therefore, based on the observed data, we examined the fit of a model with two periods of change (pre- and post-puberty) and three periods of change (up to three years before puberty, from three years before puberty to pubertal onset and from puberty to the end of follow-up) in females and males separately. The model with the best fit in females across each quartile of pubertal age was a two-spline model allowing for a single pre-pubertal change period and a single post-pubertal change period. The model with the best fit in males was a three-spline model allowing for two pre-pubertal change periods and one post-pubertal period of change.

The model for females took the form of log fat mass_ij_ = β_0_ + u_0j_ + (β_1_+ u_1j_ )s_ij1_ + (β_2_+ u_2j_ )s_ij2_ + β_3_ (age&sex adjusted height covariate)_ij_ + e_ij_ where for person j at measurement occasion i; β_0_ represents the fixed effect coefficient for the average intercept, β_1_ represents fixed effect coefficients for the average linear slope before puberty, β_2_ represents fixed effect coefficients for the average linear slope after puberty, β_3_ represents the fixed effect coefficient for the average difference in measurements between individuals of different heights, u_0j_ to u_3j_ indicate person-specific random effects for the intercept and slopes respectively, and e_ij_ represents the occasion-specific residuals or measurement error which was allowed to vary with age.

The model for males took the form of: log fat mass_ij_ = β_0_ + u_0j_ + (β_1_+ u_1j_ )s_ij1_ + (β_2_+ u_2j_ )s_ij2_ + (β_3_ + u_3j )_s_ij3_ + β_4_ (age&sex adjusted height covariate)_ij_ + e_ij_ where for person j at measurement occasion i; β_0_ represents the fixed effect coefficient for the average intercept, β_1_ represent fixed effect coefficients for the average linear slope from the first available measure to three years before puberty, β_2_ represents the fixed effect coefficients for the average linear slope from three years before puberty to pubertal onset, β_3_ represents the fixed effect coefficients for the average linear slope from pubertal onset to the end of follow-up, β_4_ represents the fixed effect coefficient for the average difference in measurements between individuals of different heights, u_0j_ to u_3j_ indicate person-specific random effects for the intercept and slopes respectively, and e_ij_ represents the occasion-specific residuals or measurement error which was allowed to vary with age

**Measurement of confounders adjusted included in main analyses**

Birthweight was extracted from medical records. Gestational age at birth was estimated from clinical records. A questionnaire at 32 weeks gestation asked mothers to report their educational attainment, which was categorized as below O-Level (Ordinary Level; exams taken in different subjects usually at age 15-16 at the completion of legally required school attendance, equivalent to today’s UK General Certificate of Secondary Education), O-Level only, A-Level (Advanced-Level; exams taken in different subjects usually at age 18), or university degree or above. Parity was defined as the number of previous pregnancies that had resulted in a live- or still-born infant collected at 18 weeks gestation. Smoking in the first trimester of pregnancy was self-reported by mothers at 18 weeks gestation; responses to smoking any tobacco (cigarettes, cigars, pipes, or other) were grouped as follows: no smoking, <10 per day, 10-19 per day or greater than 19 per day. Maternal age was reported in the mother’s antenatal questionnaires. Maternal height and weight were self-reported from the questionnaire administered at 12 weeks gestation; these were used to calculate maternal BMI. Household social class was measured as the highest of the mother’s or her partner’s occupational social class using data on job title and details of occupation collected about the mother and her partner from the mother’s questionnaire at 32 weeks gestation. Social class was derived using the standard occupational classification (SOC) codes developed by the United Kingdom Office of Population Census and Surveys and classified as I professional, II managerial and technical, IIINM non-manual, IIIM manual, and IV&V part skilled occupations and unskilled occupations. Marital status was obtained from antenatal questionnaires and classified as never married, widowed, divorced, separated, first marriage, marriage two or three. A questionnaire at 32 weeks gestation asked partners to report their educational attainment, which was categorized as below O-Level (Ordinary Level; exams taken in different subjects usually at age 15-16y at the completion of legally required school attendance, equivalent to today’s UK General Certificate of Secondary Education), O-Level only, A-Level (Advanced-Level; exams taken in different subjects usually at age 18), or university degree or above. Breastfeeding information used here was collected via questionnaires administered at 4 weeks, 6 months and 15 months.

**Table 2 Model details for log fat mass trajectories modelled using chronological age, by sex and sex-specific quartiles of pubertal age**

|  | No of contributing individuals | | Assessment of model fit | | | | |
| --- | --- | --- | --- | --- | --- | --- | --- |
|  | Total number of observations | Number of individuals with 1 measure | Mean observed (SD), log fat mass in kg | Mean predicted (SD), log fat mass in kg | Mean difference (observed – predicted), log fat mass in kg | 95% level of agreement between observed and predicted, log fat mass in kg | |
| Females |  |  |  |  |  | |  |
| Overall | 9565 | 2186 |  |  |  | |  |
| 1^st^ quartile (9.1-11.2) | 2394 | 555 | 2.78 (0.48) | 2.78 (0.46) | 0.003 | | -0.17 to 0.18 |
| 2^nd^ quartile (11.2-11.7) | 2390 | 549 | 2.65 (0.51) | 2.65 (0.49) | 0.001 | | -0.17 to 0.17 |
| 3^rd^ quartile (11.7-12.3) | 2390 | 549 | 2.56 (0.53) | 2.56 (0.51) | 0.000 | | -0.17 to 0.17 |
| 4^th^ quartile (12.3-14.6) | 2391 | 533 | 2.39 (0.56) | 2.39 (0.53) | -0.004 | | -0.18 to 0.17 |
| Males |  |  |  |  |  | |  |
| Overall | 8667 | 1990 |  |  |  | |  |
| 1^st^ quartile (10.8-13) | 2167 | 501 | 2.29 (0.62) | 2.29 (0.57) | 0.003 | | -0.28 to 0.29 |
| 2^nd^ quartile (13-13.6) | 2169 | 497 | 2.14 (0.61) | 2.14 (0.56) | 0.000 | | -0.28 to 0.28 |
| 3^rd^ quartile (13.6-14.2) | 2168 | 499 | 2.11 (0.62) | 2.11 (0.57) | 0.000 | | -0.27 to 0.27 |
| 4^th^ quartile (14.2-17.1) | 2163 | 493 | 1.96 (0.63) | 1.96 (0.58) | -0.003 | | -0.26 to 0.25 |

**Table 3 Model details for log fat mass trajectories modelled using pubertal age, by sex and sex-specific quartiles of pubertal age**

|  | No of contributing individuals | | Assessment of model fit | | | | |
| --- | --- | --- | --- | --- | --- | --- | --- |
|  | Total number of observations | Number of individuals with 1 measure | Mean observed (SD), log fat mass in kg | Mean predicted (SD), log fat mass in kg | Mean difference (observed – predicted), log fat mass in kg | 95% level of agreement between observed and predicted, ln log fat mass in kg | |
| Females |  |  |  |  |  | |  |
| Overall | 9565 | 2186 |  |  |  | |  |
| 1^st^ quartile (9.1-10.6) | 2394 | 555 | 2.78 (0.48) | 2.78 (0.44) | 0.001 | | -0.24 to 0.24 |
| 2^nd^ quartile (10.6-11.6) | 2390 | 549 | 2.65 (0.51) | 2.65 (0.47) | 0.000 | | -0.24 to 0.24 |
| 3^rd^ quartile (11.6-11.9) | 2390 | 549 | 2.56 (0.53) | 2.56 (0.49) | 0.000 | | -0.23 to 0.23 |
| 4^th^ quartile (11.9-15.7) | 2391 | 533 | 2.39 (0.56) | 2.39 (0.53) | -0.001 | | -0.24 to 0.23 |
| Males | 8667 | 1990 |  |  |  | |  |
| Overall | 2167 | 501 |  |  |  | |  |
| 1^st^ quartile (9.6-12.9) | 2169 | 497 | 2.29 (0.62) | 2.29 (0.56) | 0.001 | | -0.34 to 0.34 |
| 2^nd^ quartile (12.9-13.8) | 2168 | 499 | 2.14 (0.61) | 2.14 (0.55) | -0.001 | | -0.33 to 0.33 |
| 3^rd^ quartile (13.8-14) | 2163 | 493 | 2.11 (0.62) | 2.11 (0.56) | 0.001 | | -0.31 to 0.31 |
| 4^th^ quartile (14-18.5) | 9565 | 2186 | 1.96 (0.63) | 1.96 (0.58) | -0.001 | | -0.29 to 0.29 |

**Table 4 Results from likelihood ratio test examining linearity of association between age at peak height velocity and log fat mass at each age by sex**

|  | **Females** | **Males** |
| --- | --- | --- |
|  | **P value comparing models** | **P value comparing models** |
| Age 9 | 0.12 | 0.38 |
| Age 11 | 0.49 | 0.25 |
| Age 13 | 0.01 | 0.19 |
| Age 15 | 0.01 | 0.12 |
| Age 18 | 0.63 | 0.89 |

P value from likelihood ratio test comparing fit of a models regressing thirds of pubertal age treated as a continuous exposure on fat mass at each age to models regressing thirds of pubertal age treated as categorical exposure on fat mass at each age. P>0.05 indicates the more parsimonious model (pubertal age treated as continuous exposure) is a better fit, suggesting linearity of associations of age at peak height velocity and fat mass. Note that although the p value for females at age 13 and 15 indicated some departure from linearity, associations were still broadly linear. Thus, given the linearity of all other associations in females and males, age at peak height velocity was examined as a continuous exposure in our analyses.

**Table 5 Characteristics at birth of the mothers of children included in models compared with those excluded due to missing exposure, outcome or co-variate data**

|  | **Participants included**  **n= 4,176^a^** | **Participants excluded**  **n=1,392-15,169 ^b^** | ***P* value for comparison** † |
| --- | --- | --- | --- |
|  | **n (%)** | **n (%)** |  |
| **Maternal marital status** |  |  |  |
| Never married | 465(11.1) | 2134(22.7) | <0.001 |
| Widowed | 4(0.1) | 14(0.1) |  |
| Divorced | 135(3.2) | 444(4.7) |  |
| Separated | 35(0.8) | 184(2.0) |  |
| 1^st^ Marriage | 3265(78.2) | 5999(63.9) |  |
| Marriage 2 or 3 | 272(6.5) | 613(6.5) |  |
| **Household social class** |  |  |  |
| Professional | 767(18.4) | 774(10.4) | <0.001 |
| Managerial & Technical | 1956(46.8) | 2883(38.9) |  |
| Non-Manual | 975(23.3) | 1973(26.6) |  |
| Manual | 340(8.1) | 1224(16.5) |  |
| Part Skilled & Unskilled | 138(3.3) | 555(7.5) |  |
| **Maternal education** |  |  |  |
| Less than O level | 671(16.1) | 3085(37.1) | <0.001 |
| O level | 1475(35.3) | 2854(34.3) |  |
| A level | 1223(29.3) | 1581(19.0) |  |
| Degree or above | 807(19.3) | 803(9.6) |  |
| **Partners highest educational qualification** |  |  |  |
| Less than O level | 979(23.4) | 3175(40.5) | <0.001 |
| O level | 915(21.9) | 1641(20.9) |  |
| A level | 1223(29.3) | 1900(24.2) |  |
| Degree or Above | 1059(25.4) | 1125(14.3) |  |
| **Maternal smoking during pregnancy** |  |  |  |
| Yes | 3581(85.8) | 6416(69.8) | <0.001 |
| No | 595(14.2) | 2772(30.2) |  |
| **Parity** |  |  |  |
| 0 | 2049(49.1) | 3826(42.7) |  |
| 1 | 1480(35.4) | 3108(34.7) |  |
| 2 | 647(15.5) | 2019(22.6) |  |
| **Sex** |  |  |  |
| Female | 2186(52.3) | 7383(47.2) | <0.001 |
| Male | 2394(51.6) | 7175(47.3) |  |
|  | ***Mean (SD)*** | ***Mean (SD)*** | ***P* value** |
| Child gestational age at birth | 39.5(1.7) | 38(6.5) | <0.001 |
| Birthweight (g) | 3435.1(530.6) | 3358(599.7) | <0.001 |
| Maternal BMI (kg/m^2^) | 22.8(3.7) | 23(3.9) | 0.1 |
| Maternal age (years) | 29.5(4.4) | 27(5.1) | <0.001 |
| Mean age at peak height velocity (years) | 12. 6 (1.3) | 12.6 (1.3) | 0.17 |

^a^ Denominators for excluded participants in this table varies due to missing data for characteristics shown.

**Table 6 Unadjusted mean trajectory and mean difference in trajectory of height-adjusted fat mass per year later age at peak height velocity, from chronological age multilevel models**

|  | **Mean trajectory (95% CI) of height-adjusted fat mass** | | **Mean difference in height-adjusted fat mass (95% CI)** | |
| --- | --- | --- | --- | --- |
|  | Age 9y (kg) ^a^ | 7.40 (1.54) | Age 9y (% difference) ^c^ | -21.24 (-23.16,-19.33) |
|  | 9-13y (% /y) ^b^ | 16.40 (3.71) | 9-13y (% difference /y) ^d^ | 0.50 (0.04,0.97) |
| **Females** | 13-15y (%/y) ^b^ | 11.11 (7.26) | 13-15y (% difference /y) ^d^ | 2.72 (1.90,3.54) |
|  | 15-18y (% /y) ^b^ | 6.05 (3.83) | 15-18y (% difference /y) ^d^ | 2.39 (1.84,2.95) |
|  | Age 18y (kg) ^a^ | 20.01 (1.04) | Age 18y (% difference) ^c^ | -8.98 (-10.9,-7.05) |
|  | Age 9y (kg) ^a^ | 5.36 (1.60) | Age 9y (% difference) ^c^ | -23.31 (-25.40,-21.22) |
|  | 9-13y (% /y) ^b^ | 16.37 (3.41) | 9-13y (% difference /y) ^d^ | 4.69 (4.09,5.29) |
| **Males** | 13-15y (%/y) ^b^ | -7.24 (8.32) | 13-15y (% difference /y) ^d^ | 0.96 (-0.11,2.03) |
|  | 15-18y (% /y) ^b^ | 10.32 (6.10) | 15-18y (% difference /y) ^d^ | -2.51 (-3.34,-1.68) |
|  | Age 18y (kg) ^a^ | 11.36 (1.75) | Age 18y (% difference) ^c^ | -12.98 (-15.85,-10.11) |

Mean trajectory is centred on the sex-specific mean of age at peak height velocity for each sex (age ~11.7 for females and age ~13.6 for males).The difference in fat mass per year of age at peak height velocity is back transformed from the log scale for ease of interpretation and is a ratio of geometric means, expressed as a percentage difference.

^a^ Mean height-adjusted fat mass at 9y and 18y in kg. ^b^ Percentage change per year in height-adjusted fat mass. ^c^  Percentage difference in fat mass at 9y and 18y per year later age at peak height velocity. ^d^ Percentage difference in change per year, per year later age at peak height velocity.

CI, confidence interval.

**Table 7 Unadjusted mean trajectory and mean difference in trajectory of height-adjusted fat mass per year later age at peak height velocity, from pubertal age multilevel models**

|  | **Mean trajectory (95% CI) of height-adjusted fat mass** | | **Mean difference in height-adjusted fat mass (95% CI)** | |
| --- | --- | --- | --- | --- |
|  | Before puberty (% /y) ^a^ | 19.4 (2.50) | Before puberty (% difference /yr) ^c^ | -1.69 (-2.45,-0.92) |
| **Females** | Fat mass at puberty (kg) ^b^ | 11.90 (1.50) | Fat mass at puberty (% difference) ^d^ | -10.18 (-12.20,-8.16) |
|  | After puberty (%/y) ^a^ | 9.61 (3.62) | After puberty (% difference /yr) ^c^ | 1.44 (1.11,1.76) |
|  | Up-to 3 years before puberty (% /y) ^a^ | 26.68 (2.96) | Up-to 3 years before puberty (% difference/y ) ^c^ | -5.84 (-7.17,-4.5) |
| **Males** | From 3 years before to puberty (% /y) ^a^ | 2.47 (5.47) | From 3 years before to puberty (% difference/yr) ^c^ | 3.08 (2.19,3.96) |
|  | Fat mass at puberty (kg)  ^b^ | 8.59 (1.76) | Fat mass at puberty (% difference) ^d^ | -3.57 (-6.49,-0.66) |
|  | After puberty (% /y ) ^a^ | 3.79 (4.09) | After puberty (% difference /y ) ^c^ | -0.75 (-1.36,-0.14) |

Mean trajectory is centred on the sex-specific mean of age at peak height velocity for each sex (age ~11.7 for females and age ~13.6 for males).The difference in fat mass per year of age at peak height velocity is back transformed from the log scale for ease of interpretation and is a ratio of geometric means, expressed as a percentage difference.

^a^ Mean height-adjusted fat mass at 9y and 18y in kg. ^b^ Percentage change per year in height-adjusted fat mass. ^c^  Percentage difference in fat mass at 9y and 18y per year later age at peak height velocity. ^d^ Percentage difference in change per year, per year later age at peak height velocity.

CI, confidence interval.

|  | **Association of age at peak height velocity with fat mass from regression**  **(95% CI)** | **Association of age at peak height velocity with fat mass from multilevel model based on pubertal age**  **(95% CI)** | **Association of age at peak height velocity with fat mass from multilevel model based on chronological age**  **(95% CI)** |
| --- | --- | --- | --- |
| **Females** |  |  |  |
| **Log fat mass (kg)** ^a^ |  |  |  |
| Age 9 | 0.24 (0.22, 0.27) | 0.22 (0.19,0.25) | 0.24 (0.22,0.26) |
| Age 18 | 0.11 (0.09, 0.14) | 0.13(0.10, 0.15) | 0.09 (0.07,0.11) |
| **Males** |  |  |  |
| **Log fat mass (kg)** ^a^ |  |  |  |
| Age 9 | 0.21 (0.18, 0.24) | 0.32 (0.28,0.35) | 0.28 (0.25,0.31) |
| Age 18 | 0.13 (0.09, 0.17) | 0.10 (0.07,0.13) | 0.14 (0.11,0.18) |

Table 8 Association of pubertal timing with first and last available measure of fat mass using linear regression compared with predicted differences at 9 and 18 years from multilevel models

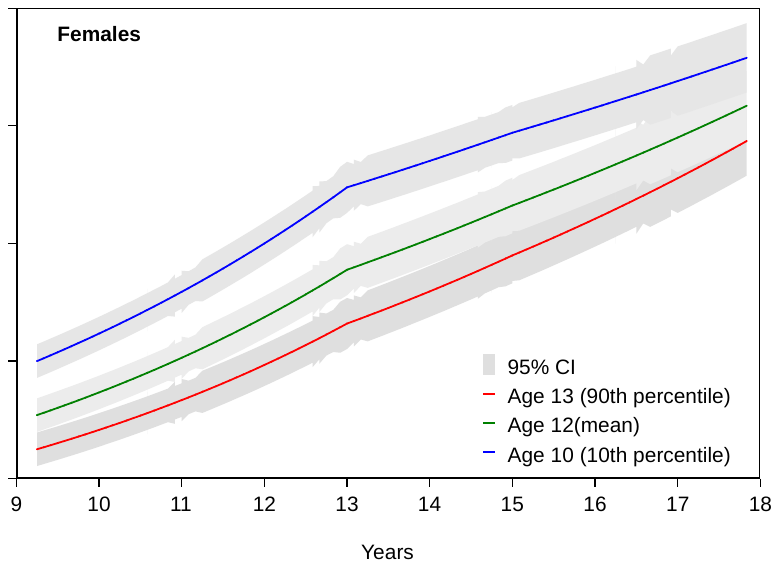

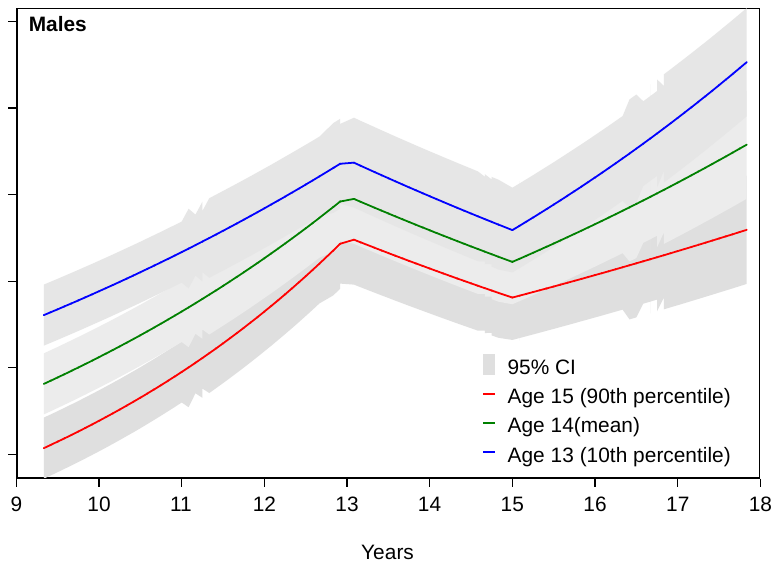

Figure 2 Mean predicted trajectories of fat mass by pubertal age in females and males from models based on chronological age, restricted to participants with at least one measure before and one measure after puberty

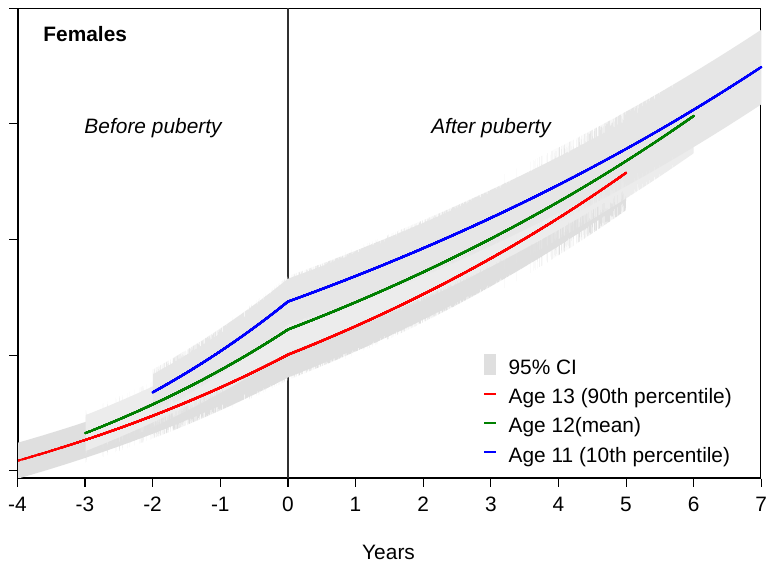

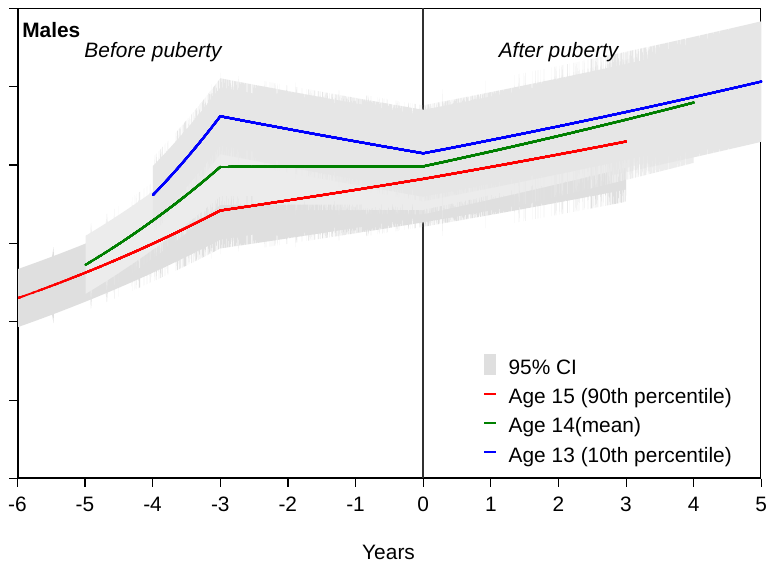

**Figure 3 Mean predicted trajectories of fat mass by pubertal age in females and males from models based on pubertal age, restricted to participants with at least one measure before and one measure after puberty**

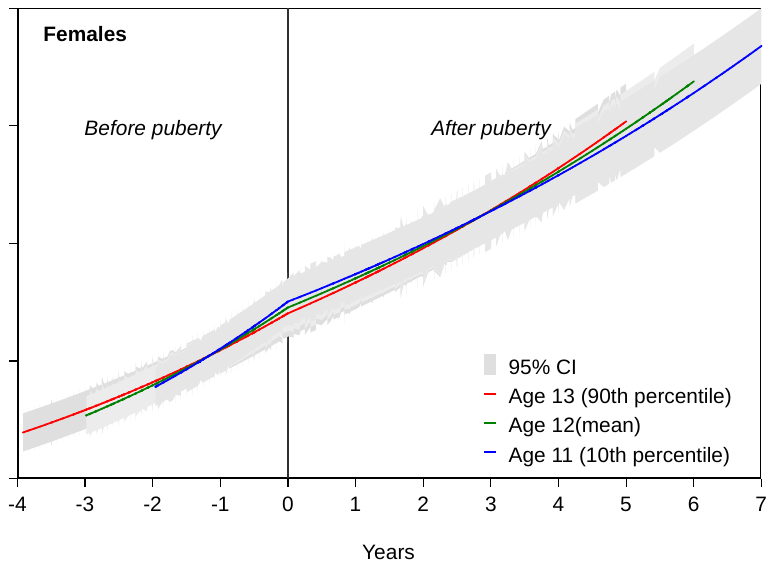

**Figure 4 Mean predicted trajectories of fat mass by pubertal age (based on age at menarche) in females from models based on pubertal age**

1. Frysz M, Howe LD, Tobias JH, et al. Using SITAR (SuperImposition by Translation and Rotation) to estimate age at peak height velocity in Avon Longitudinal Study of Parents and Children. *Wellcome Open Research* 2018;3

2. Cole TJ, Donaldson MD, Ben-Shlomo Y. SITAR--a useful instrument for growth curve analysis. *International journal of epidemiology* 2010;39(6):1558-66. doi: 10.1093/ije/dyq115 [published Online First: 2010/07/22]

3. Simpkin AJ, Sayers A, Gilthorpe MS, et al. Modelling height in adolescence: a comparison of methods for estimating the age at peak height velocity. *Annals of human biology* 2017;44(8):715-22.

4. O'Keeffe L, Simpkin A, Tilling K, et al. Sex-specific trajectories of cardiometabolic risk factors during childhood and adolescence: a prospective cohort study Atherosclerosis. *Atherosclerosis* 2018;278:190-96.

5. O'Keeffe L, Simpkin A, Tilling K, et al. Data on trajectories of measures of cardiovascular health in the Avon Longitudinal Study of Parents and Children (ALSPAC). *Data in Brief* 2018
